# Histone neutralization in a rat model of acute lung injury induced by double-hit lipopolysaccharide

**DOI:** 10.1101/2021.12.26.474211

**Authors:** Yangyang Ge, Chenchen Wang, Yuduo Zhen, Junjie Luo, Jiayi Chen, Yu Wang, Fuquan Wang, Li Wang, Yun Lin, Lin Shi, Shanglong Yao

## Abstract

**Background:** Acute respiratory distress syndrome (ARDS) remains a challenge because of its high morbidity and mortality. Circulation histones levels in ARDS patients were correlated to disease severity and mortality. This study examined the impact of histone neutralization in a rat model of acute lung injury (ALI) induced by a lipopolysaccharide (LPS) double-hit.

**Methods:** Sixty-eight male Sprague-Dawley rats were randomized to sham (N=8, received saline only) or LPS (N=60). The LPS double-hit consisted of a 0.8 mg/kg intraperitoneal injection followed after 16 hours by 5 mg/kg intra-tracheal nebulized LPS. The LPS group was then randomized into five groups: LPS only (N=12); LPS + 5, 25, or 100 mg/kg intravenous STC3141 every 8 hours (LPS+L, LPS+M, LPS+H, respectively, each N=12); or LPS + intraperitoneal dexamethasone 2.5 mg/kg every 24 hours for 56 hours (LPS+D, N=12) The animals were observed for 72 hours.

**Results:** LPS animals developed ALI as suggested by lower oxygenation, lung edema formation, and histological changes compared to the sham animals. Compared to the LPS group, LPS+H and +D animals had significantly lower circulating histone levels; only the LPS+D group had significantly lower bronchoalveolar lavage fluid (BALF) histone concentrations. The LPS+L, +M, +H and +D groups had improved oxygenation compared to the LPS group and the LPS+H and +D groups had a lower lung wet-to-dry ratio. All animals survived.

**Conclusion:** Neutralization of histone using STC3141, especially at high dose, had similar therapeutic effects to dexamethasone in this LPS double-hit rat ALI model, with significantly decreased circulating histone concentration, improved oxygenation, and decreased lung edema formation.

## I. Introduction

Acute respiratory distress syndrome (ARDS), characterized by bilateral chest radiographical opacities and severe hypoxemia (1), remains a major challenge with its high morbidity and mortality. In an observational study across 50 countries and involving 459 intensive care units (ICUs), ARDS was responsible for 10% of ICU admissions and 23% of mechanically ventilated patients; mortality was 35% and could be as high as 46% in severe ARDS (2). Currently, only physiological manipulation, such as low tidal volume and prone position ventilation, has been shown to have a survival benefit in these patients, and there are no pharmaceutical therapies available (3).

Neutrophils are the first line immune cells against microorganisms, acting by phagocytosis, release of reactive oxygen species (ROS), degranulation (4) and decondensed chromatin fibers coated with antimicrobial proteins, such as histones and neutrophil elastase, which form neutrophil extracellular traps (NETs) to trap and kill bacteria (5). However, NETs may be double-edged swords (6), with excessive formation playing an important role in the pathogenesis of ARDS (7): NETs expand into the pulmonary alveoli, causing lung injury, and induce epithelial and endothelial cell death (7). NETs also provoke the formation of immuno-thrombosis (8), which is associated with worse ARDS outcomes. There is therefore a sound rationale for strategies that target histones and decrease NET formation in the treatment of ARDS.

Recent studies showed that extracellular histones significantly increased the development of ARDS. In an observational study that included 96 patients with ARDS and 30 healthy volunteers, extracellular histone levels from plasma and bronchoalveolar fluid (BALF) were highly correlated to the severity of ARDS and were significantly higher in non-survivors (9). In 52 patients with non-thoracic trauma, high circulating histone levels were associated with an increased incidence of acute lung injury (ALI) and higher Sequential Organ Failure Assessment (SOFA) scores (10). Interestingly, anti-histone antibody therapy protected mice from histone-induced lethality (10). Targeting extracellular histones may thus represent a promising therapeutic option for ARDS (11).

A polyanion molecule of beta-O-methyl cellobiose sulfate sodium salt (mCBS.Na), STC3141, has been identified as able to neutralize histones and decrease histone-induced red blood cell (RBC) and platelet aggregation in vitro (12). Its use was associated with a survival benefit in a rat cecal ligation and puncture model (12). This molecule has also been shown to decrease infarct size in a rat cardiac ischemia/reperfusion model (13). Beneficial effects in ARDS are therefore anticipated.

Our hypothesis was that mCBS.Na (STC3141) administration would improve outcomes in a rat model of ALI induced by a lipopolysaccharide (LPS) double-hit. Dexamethasone administration has been shown to increase ventilator-free days and decrease 60-day mortality in patients with ARDS (14), so we decided to use dexamethasone as a positive control in this study. The reason we chosen this rat model is that it closely reproduces the acute phase of human ARDS (15-17).

## II. Materials and methods

### 2.1 ALI rat model and experimental protocol

All procedures in this study were conducted in accordance with the guidelines of the National Institute of Health and approved by the Animal Protection and Utilization Committee of the JOINN laboratory.

Six- to eight-week-old male Sprague-Dawley (SD) rats (195g-270g) (Vital River Laboratory Animal Technology Co., Ltd. Zhejiang) were raised in a pathogen-free environment at temperature 20-26°C under a natural light and dark cycle with free access to standard chow and water. A double-hit LPS strategy (LPS, Sigma-Aldrich, St. Louis, MO, USA) was used to create the model.

A total of 68 adult male SD rats were randomized to a sham group (Sham, N= 8), in which animals received intraperitoneal injection and intra-tracheal nebulized saline only, or a double-hit LPS group (N=60). The double-hit LPS consisted of 0.8 mg/kg intraperitoneal injection followed after 16 hours by 5 mg/kg intra-tracheal nebulized LPS. For the nebulized LPS or saline, animals were anesthetized by inhalation of isoflurane, then fixed on a 45° inclined rat holder plate. An anesthetic laryngoscope was used to pin the animal’s tongue and expose the glottis. A micro liquid atomizer loaded with LPS solution or saline was inserted into the trachea and the solution injected quickly into the lungs. The rats were then placed vertically and rotated for 0.5-1 min to insure a uniform distribution of solution in the lung.

The LPS animals were then randomized to an ARDS control group (LPS, n=12), in which animals received an intravenous injection of saline every 8 hours for 64 hours; a LPS+L group (n=12), animals received an intravenous injection of 5 mg/kg STC3141 every 8 hours for 64 hours; LPS+M group (n=12), animals received an intravenous injection of 25 mg/kg STC3141 every 8 hours for 64 hours; LPS+H group (n=12), animals received an intravenous injection of 100 mg/kg STC3141 every 8 hours for 64 hours; or a positive control group (ARDS+D, n=12) in which animals received an intra-peritoneal injection of 2.5 mg/kg dexamethasone (KingYork, Tianjin, China) every 24 hours for a total of 56 hours (Figure1).

**Figure1.**
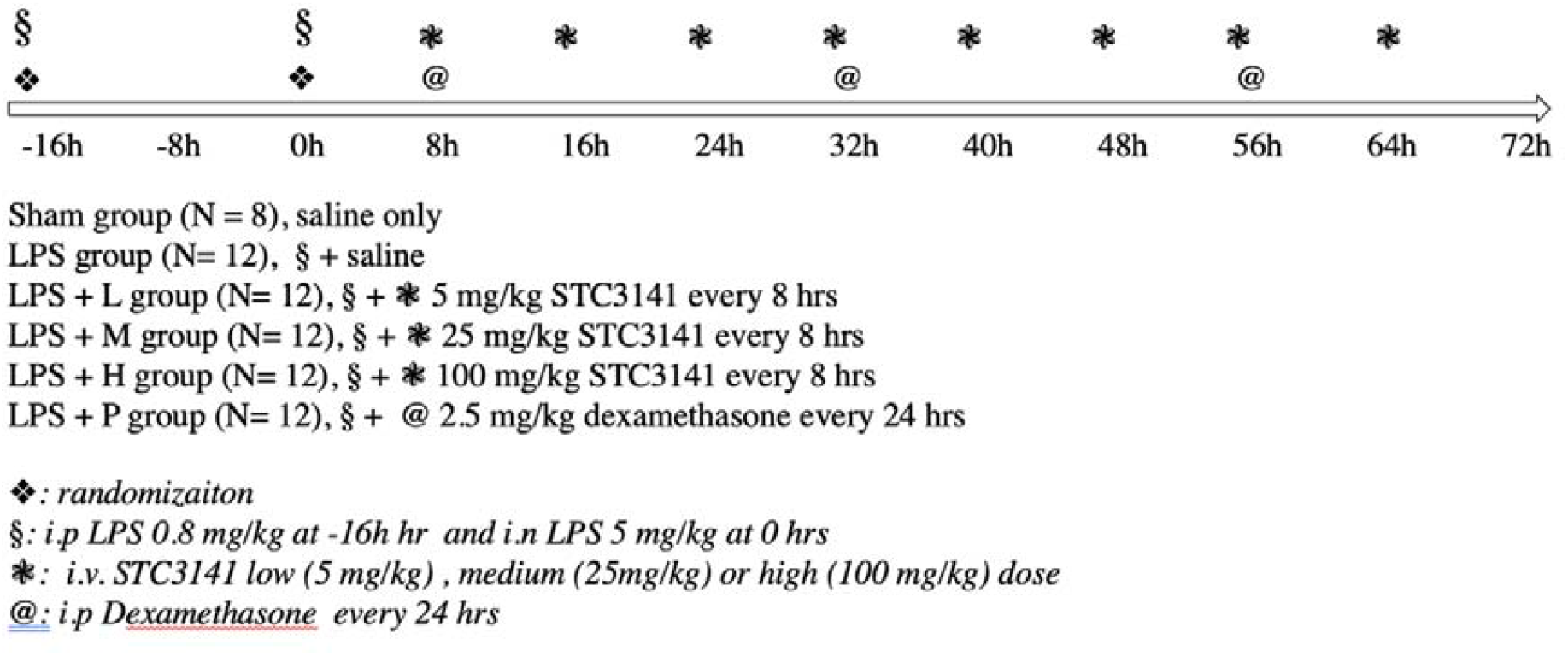
Experimental protocol

### 2.2 Euthanasia, autopsy, blood sampling, and BALF collection

In accordance with the American Veterinary Medical Association Guidelines (2013), at 72 hours after intra-tracheal LPS nebulization, rats were anesthetized with chloral hydrate (350 mL/kg, 100 mg/mL), and then euthanized after collection of 0.5 ml arterial blood from the abdominal aorta, which was exposed by opening the abdomen along a medioventral line.

To collect BALF, a tracheal cannula was slowly inserted into the trachea through an incision in the left bronchus and fixed in the centripetal direction with threads. A total volume of 3 mL PBS buffer was slowly injected to fill the lung and thereafter gently withdrawn. This procedure was repeated 3 times, for 10 s each time, and the BALF was collected into a centrifuge tube. It was then centrifuged at 4°C with 1200 g for 10 minutes. The supernatant was separated and stored at -80 °C. The precipitation was resuspended with 1 mL PBS buffer for cell counting. Cell counting and classification of leukocytes in the BALF was performed using an automatic blood cell analyzer.

The middle lobe of the right lung was isolated and weighed to calculate the wet-to-dry ratio. The remaining right lung tissue and bronchi were fixed in 10% neutral buffered formalin solution, paraffin-embedded, sectioned, and hematoxylin-eosin (HE) stained for pathomorphological examination of the lung tissue.

### 2.3 Blood gas and histone measurements in plasma and BALF

At autopsy, a blood gas analyzer (i-STAT ®1 Handheld Blood Analyzer (300-G. Abbott Point of Care Inc. USA)) was used to measure PO2 (mmHg), PCO2 (mmHg), pH, and SO2%. Blood samples were collected and centrifuged at 4°C for about 3000 rpm. The plasma was collected and stored at -70°C. Histone concentration was measured using an ELISA kit (Roche, Sigma-Aldrich, St. Louis, MO, USA) according to the producer’s instructions.

### 2.4 Histology and inflammation

Pathological morphological examination, diagnosis, and classification were conducted according to a four-grade method (minimal, slight, moderate, severe).

### 2.5 Lung wet-to-dry weight

The right middle lobe was weighed, then heated to 60°C for 72 h and weighed again. The wet-to-dry weight ratio was calculated using the formula:lung wet-to-dry ratio = right middle lobe wet weight/dry weight.

### 2.6 Data collection and statistical analysis

We used Prism 9.3.0 (345) statistical software to process the data and draw the figures. Statistical analysis was performed using a one-way ANOVA with Tukey test. All data are expressed as “mean ± SD”. All statistical tests were conducted as 2-sided tests, and the level of significance was set at P < 0.05.

## III. Results

The animals in the LPS group developed acute lung injury, with arterial blood PO2 (77.5 ± 11.4 mmHg versus 103.3 ± 5.9 mmHg) and oxygen saturation (95.3 ± 2.5% versus 98.4 ± 0.5%) significantly lower in the LPS group than in the sham group (Table 1). The WBC (9.41 ± 2.01 vs 0.32 ± 0.22) and neutrophil (5.17 ± 1.90 vs 0.04 ± 0.02) counts in the BALF, and the BALF histone concentration (0.835 ± 0.380 vs 0.039 ± 0.012) were significantly higher in the LPS group than in the sham group. The lung wet-to-dry ratio was significantly higher in the LPS group (2.26 ± 0.51 vs 1.17 ± 0.09).

**Table 1:**
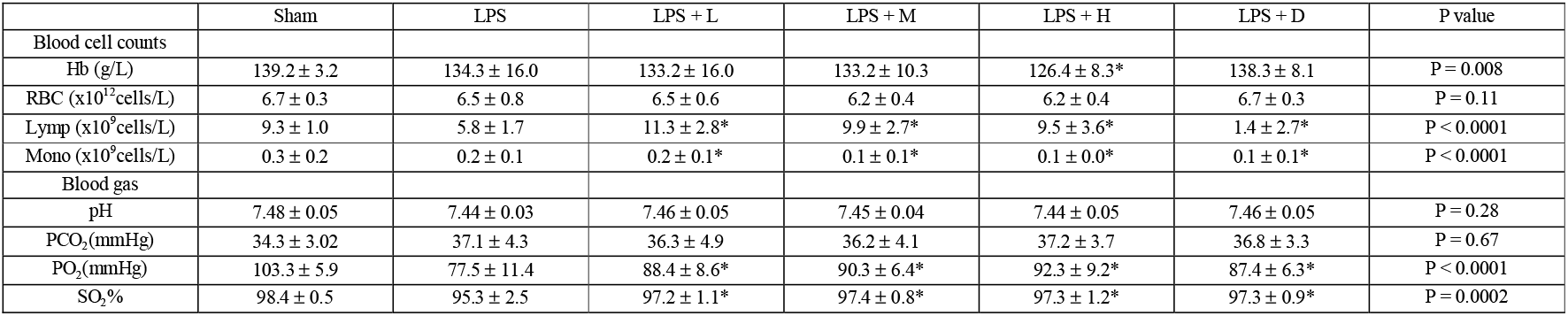
Hemoglobin (Hb) concentration and red blood cell (RBC), lymphocyte (Lymp) and monocyte (Mono) counts in the blood, arterial pH, PCO2, PO2 and SO2% in the different groups. * indicates significant difference compared with LPS group.

### Comparison between LPS groups

The arterial PO2 was significantly higher in the LPS+L, +M, and +H groups than in the LPS only group; the oxygen saturation was significantly higher in the LPS+D group than in the LPS only group. There were no statistical differences in PCO2 or pH among groups (Table 1).

WBC, lymphocyte, and platelet counts were significantly lower in the LPS+L, +M, +H and +D groups than in the LPS only group (Figure 2).

**Figure 2.**
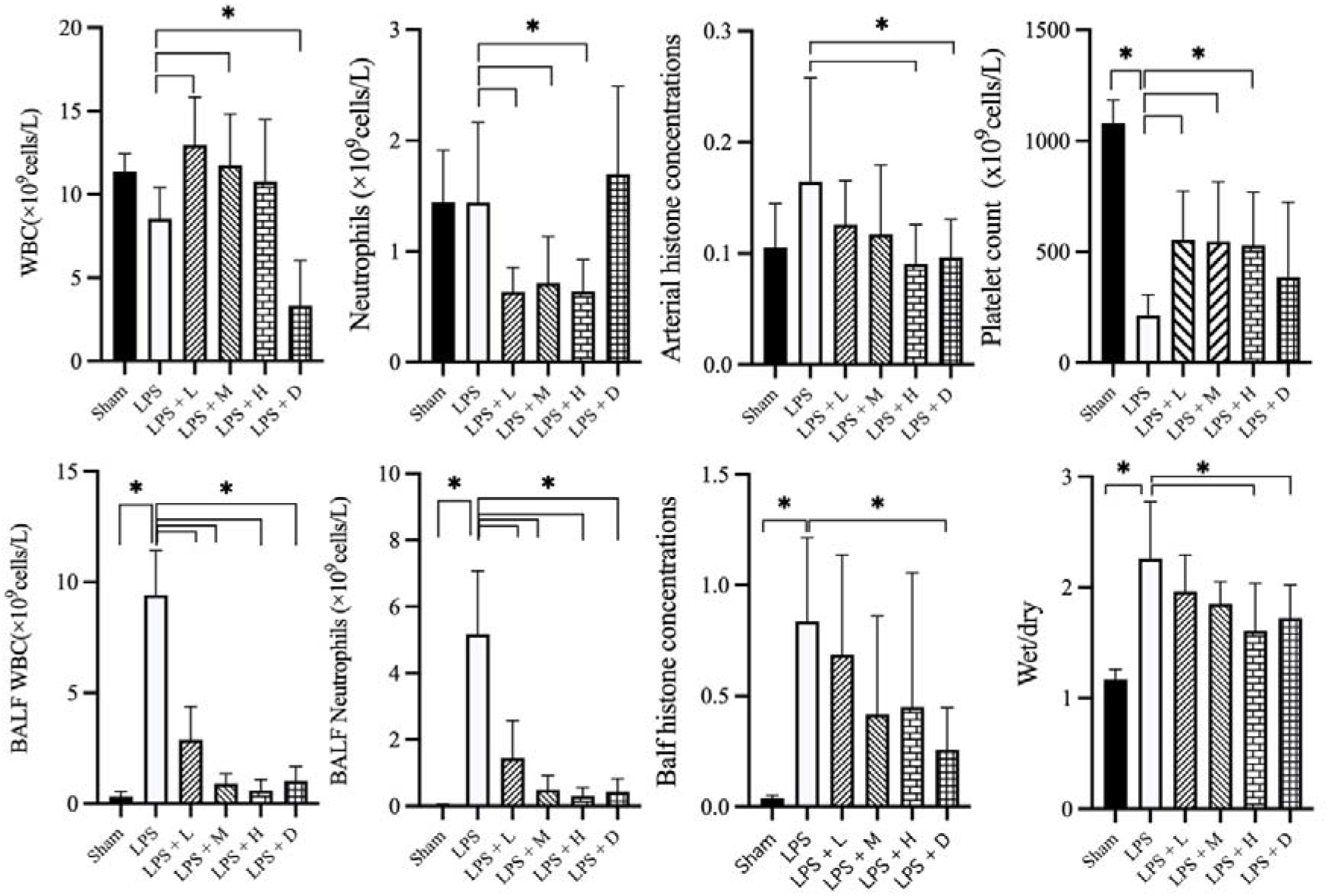
Blood and bronchoalveolar lavage fluid (BALF) white blood cell (WBC), neutrophil, and platelet counts, arterial and BALF histone concentrations, and lung wet-to-dry ratio in the different groups: data expressed as mean ± standard deviation; * stands for p<0.05.

Animals in the LPS+H and +D groups had significantly lower circulating histone levels than the other LPS groups; the LPS+D group also had lower BALF histone concentrations (Figure 2).

Histopathological examination showed lung inflammation, with neutrophil-based inflammatory cells (alveolar cavity, blood vessel, alveolar wall), alveolar wall thickening, bleeding in the alveoli (with or without hemoglobin crystals). The degree of inflammation was significantly less in the LPS+L, +M, +H and +D groups than in the LPS only group. Inflammation in the LPS+L, +M and +H animals showed a dose-dependent response: as the dose increased, the degree of severity decreased. Representative histological findings in the right upper lobe in the different groups are shown in Figure 3.

**Figure 3.**
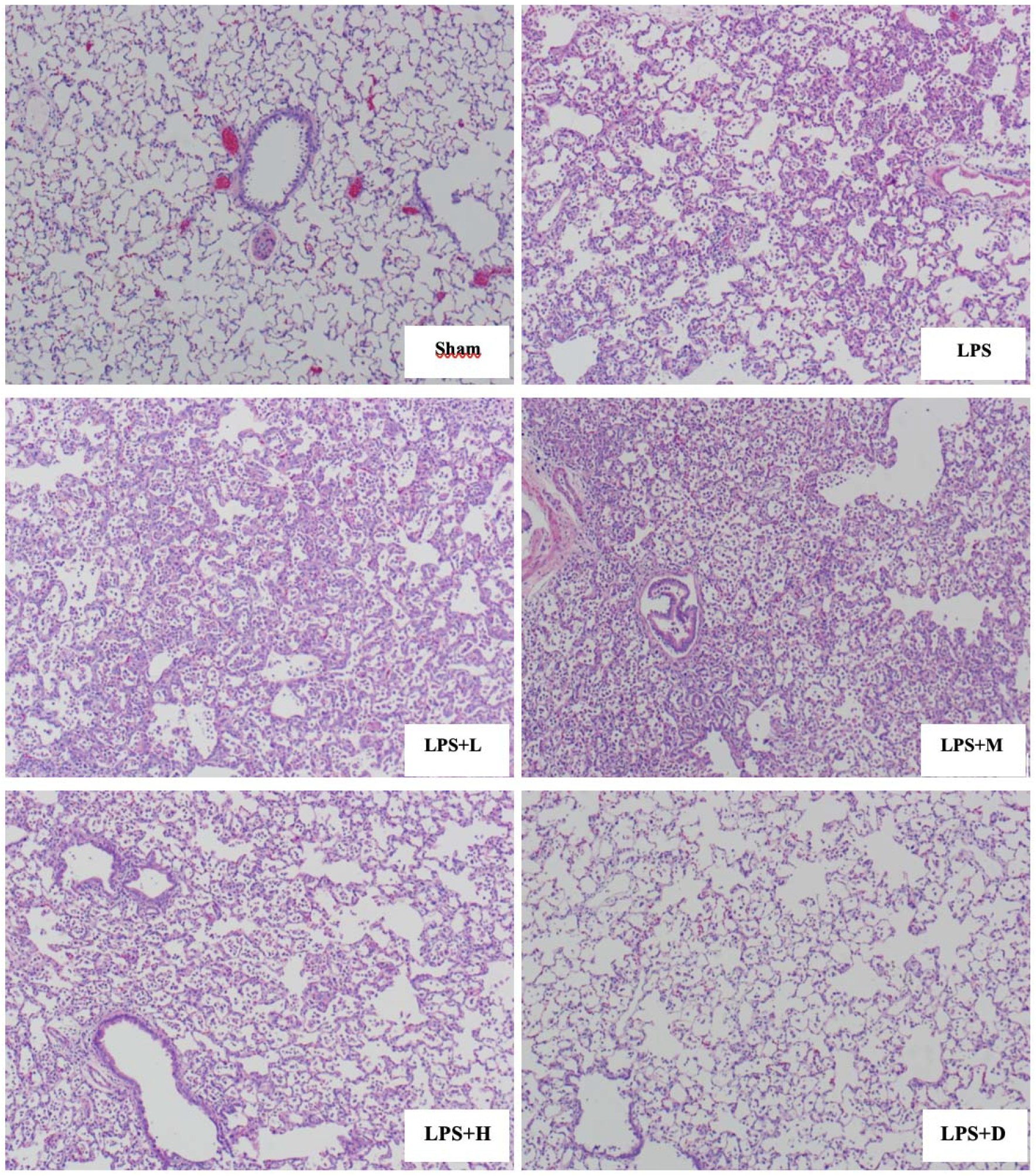
Representative histology of the right upper lobe in the different groups: in sham group: normal; LPS group, severe inflammation; LPS+L group, severe inflammation; LPS+M group severe inflammation; LPS+H group: moderate inflammation; LPS+D group: minimal inflammation. All images are taken after HE staining and amplified 100X.

The LPS+H and +D animals had a significantly lower lung wet-to-dry ratio than that of the LPS only group (Figure 2). All the 68 animals survived to 72 hours after the LPS second hit.

## IV. Discussion

The main findings of the current study are: 1) that a double-hit of LPS induced a model of ALI in rats as shown by the presence of inflammation, decreased oxygenation, lung edema, and histological changes, representing clinical ARDS findings; 2) neutralization of extracellular histone with low, medium and high dose STC3141 significantly decreasing circulating and BALF WBC and neutrophil counts, improving lung oxygenation and decreasing lung inflammation; 3) high dose STC3141 seems to have a similarly beneficial effect on oxygenation and lung edema formation to dexamethasone administration in this ALI model.

LPS exposure may lead to lung inflammation. An LPS stimulated rat model is a standard experimental ALI and ARDS model and widely used for new drug development for this condition (18-20). LPS stimulation leads to systemic inflammatory and toxic responses by activating Toll-like receptors (21), the complement system (22), and inflammasome pathways, which provoke apoptosis, necrosis, and NET formation (7), and histone release into the circulation. These extracellular histones play a pathogenic role in the development of ARDS through promoting alveolar macrophage proptosis (23), direct cytotoxic endothelial damage (24) and pro-inflammatory effects with cytokine production (25)(26), resulting in overwhelming cell damage and death (22). Furthermore, these extracellular histones can induce platelet aggregation (28) and coagulation activation (29), immuno-thrombosis formation (30) and innate immunity by activating Toll-like receptors and the NLRP3 inflammasome (27). In our study, LPS animals exhibited decreased oxygenation, lung inflammation (intra-alveolar leukocytes, histological injury), and higher histone concentrations than sham animals. STC3141 significantly decreased WBC and neutrophil counts not only in the circulation, but also in the BALF, which suggests it may interrupt NET formation and subsequent immunothrombosis, thereby ameliorating the impaired microcirculation and improving lung function.

Targeting extracellular histones using STC3141 improved oxygenation and lung edema formation in the current ARDS model: low and medium dose STC3141 treatment did not decrease circulating histone levels, but high dose STC3141 and dexamethasone did. Interestingly, in BALF, only dexamethasone decreased histone concentration, which raises the question as to whether high dose STC3141 is sufficient? An on time measurement of circulating levels of histones may help to answer this question, but this approach is currently under development. This dose-dependent effect is reflected by the findings of similar lung dry/wet ratio in the high dose and dexamethasone groups: improved oxygenation and decreased lung inflammation and edema formation. Meara et al similarly showed that mCBS.Na administration significantly decreased histone-induced RBC and platelet aggregation in vitro, and histone-induced tissue injury, thrombocytopenia and anemia in mice (12). These preclinical findings could help to explain the observations in ARDS patients, with a significant increase in plasma histones in mild, moderate, and severe ARDS and significantly higher plasma histones in non-survivors than in survivors (9). Furthermore, in ARDS patients with a good prognosis, plasma histone levels decreased after admission whereas they increased in those with a poor prognosis (9). All these findings support the concept of histone neutralization therapy in ARDS.

The study has several limitations. First, LPS is well known to induce inflammation rather than infection, and the model therefore does not fully mimic the disease kinetics in ARDS (31). Second, the young healthy animals used in the current study are different from the typical ARDS patient with comorbidities. Third, in the present ARDS model, all the rats survived, which may suggest it is a mild injury model; in addition, no hypoxemia was observed. Fourth, no antibiotics, fluid resuscitation, vasopressors, or mechanical ventilation were used, limiting the clinical relevance of the model. Fifth, we did not perform immunohistochemistry to determine NET formation in the lung. Sixth, the ambient environment temperature is 28°C for the rat (32), while our experiments processed in a relative cold environment, this cold stress response may also bias our results. Seventh, use of specific pathogen-free animals may also bias the results, because their immune response might different from wild animals(33). Eighth, the shortage of direct evidence immunohistochemistry results of NETs from lung with STC3141 due to limited label technology.

Three clinical trials are currently ongoing to test the effects of STC3141: a phase II study in patients with coronavirus disease 2019 (COVID-19), which has currently finished patients inclusion (ClinicalTrials.gov Identifier: NCT04880694); a phase Ib study in sepsis, which plans to include 25 patients (); and a phase Ib study in ARDS, which plans to include… patients ().

## V. Conclusion

In the current LPS double-hit rat model of ALI, high dose STC3141 showed similar effects to dexamethasone therapy, including significantly decreased circulating and BALF histone concentrations, improved oxygenation, and decreased lung edema formation. Clinical trials are necessary to identify the effects of histone neutralization in ARDS patients.

## Acknowledgments

No.

## VI.References

1. ARDS Definition Task Force, Ranieri VM, Rubenfeld GD, Thompson BT, Ferguson ND, Caldwell E, et al. Acute respiratory distress syndrome: the Berlin Definition. JAMA. 2012 Jun 20;307(23):2526–33.

2. Bellani G, Laffey JG, Pham T, Fan E, Brochard L, Esteban A, et al. Epidemiology, patterns of care, and mortality for patients with acute respiratory distress syndrome in intensive care units in 50 countries. JAMA. 2016 Feb 23;315(8):788–800.

3. Meyer NJ, Gattinoni L, Calfee CS. Acute respiratory distress syndrome. Lancet Lond Engl. 2021;398(10300):622–37.

4. Segal AW. How neutrophils kill microbes. Annu Rev Immunol. 2005;23:197–223.

5. Brinkmann V, Reichard U, Goosmann C, Fauler B, Uhlemann Y, Weiss DS, et al. Neutrophil extracellular traps kill bacteria. Science. 2004 Mar 5;303(5663):1532–5.

6. Kaplan MJ, Radic M. Neutrophil extracellular traps: double-edged swords of innate immunity. J Immunol Baltim Md 1950. 2012 Sep 15;189(6):2689–95.

7. Porto BN, Stein RT. Neutrophil extracellular traps in pulmonary diseases: too much of a good thing? Front Immunol. 2016 Aug 15;7:311

8. Yago T, Liu Z, Ahamed J, McEver RP. Cooperative PSGL-1 and CXCR2 signaling in neutrophils promotes deep vein thrombosis in mice. Blood. 2018 Sep 27;132(13):1426–37.

9. Lv X, Wen T, Song J, Xie D, Wu L, Jiang X, et al. Extracellular histones are clinically relevant mediators in the pathogenesis of acute respiratory distress syndrome. Respir Res. 2017;18:165

10. Abrams ST, Zhang N, Manson J, Liu T, Dart C, Baluwa F, et al. Circulating histones are mediators of trauma-associated lung injury. Am J Respir Crit Care Med. 2013 Jan 15;187(2):160–9.

11. Karki P, Birukov KG, Birukova AA. Extracellular histones in lung dysfunction: a new biomarker and therapeutic target? Pulm Circ. 2020 Dec;10(4):2045894020965357.

12. Meara CHO, Coupland LA, Kordbacheh F, Quah BJC, Chang C-W, Simon Davis DA, et al. Neutralizing the pathological effects of extracellular histones with small polyanions. Nat Commun. 2020 Dec;11(1):6408.

13. Shah M, He Z, Rauf A, Beikoghli Kalkhoran S, Heiestad CM, Stensløkken K-O, et al. Extracellular histones are a target in myocardial ischaemia–reperfusion injury. Cardiovasc Res. 2021 Apr 20;cvab139.

14. Villar J, Ferrando C, Martínez D, Ambrós A, Muñoz T, Soler JA, et al. Dexamethasone treatment for the acute respiratory distress syndrome: a multicentre, randomised controlled trial. Lancet Respir Med. 2020 Mar;8(3):267–76.

15. Brigham KL, Meyrick B. Endotoxin and lung injury. Am Rev Respir Dis. 1986 May;133(5):913–27.

16. Zhao S, Zhang Y, Chen Q, Dong S, Zhang G, Li J, et al. A modified “double-hit” induced acute lung injury model in rats and protective effects of tetramethylpyrazine on the injury via Rho/ROCK pathway. Int J Clin Exp Pathol. 2015 May 1;8(5):4581–7.

17. Chimenti L, Morales-Quinteros L, Puig F, Camprubi-Rimblas M, Guillamat-Prats R, Gómez MN, et al. Comparison of direct and indirect models of early induced acute lung injury. Intensive Care Med Exp. 2020 Dec 18;8(Suppl 1):62.

18. Martin MA, Silverman HJ. Gram-negative sepsis and the adult respiratory distress syndrome. Clin Infect Dis Off Publ Infect Dis Soc Am. 1992 Jun;14(6):1213–28.

19. Liang Q, Lin Q, Li Y, Luo W, Huang X, Jiang Y, et al. Effect of SIS3 on Extracellular Matrix Remodeling and Repair in a Lipopolysaccharide-Induced ARDS Rat Model. J Immunol Res. 2020;2020:6644687.

20. Uhlig S, Kuebler WM. Difficulties in modelling ARDS (2017 Grover Conference Series). Pulm Circ. 2018 Jun;8(2):2045894018766737.

21. Chen R, Kang R, Fan X-G, Tang D. Release and activity of histone in diseases. Cell Death Dis. 2014 Aug 14;5:e1370.

22. Xu Z, Huang Y, Mao P, Zhang J, Li Y. Sepsis and ARDS: The dark side of histones. Mediators Inflamm. 2015;2015:205054.

23. Jiang P, Jin Y, Sun M, Jiang X, Yang J, Lv X, et al. Extracellular histones aggravate inflammation in ARDS by promoting alveolar macrophage pyroptosis. Mol Immunol. 2021 Jul;135:53–61.

24. Xu J, Zhang X, Pelayo R, Monestier M, Ammollo CT, Semeraro F, et al. Extracellular histones are major mediators of death in sepsis. Nat Med. 2009 Nov;15(11):1318–21.

25. Xu J, Zhang X, Monestier M, Esmon NL, Esmon CT. Extracellular histones are mediators of death through TLR2 and TLR4 in mouse fatal liver injury. J Immunol Baltim Md 1950. 2011 Sep 1;187(5):2626–31.

26. Zhang Y, Zhao J, Guan L, Mao L, Li S, Zhao J. Histone H4 aggravates inflammatory injury through TLR4 in chlorine gas-induced acute respiratory distress syndrome. J Occup Med Toxicol Lond Engl. 2020;15:31.

27. Allam R, Kumar SVR, Darisipudi MN, Anders H-J. Extracellular histones in tissue injury and inflammation. J Mol Med Berl Ger. 2014 May;92(5):465–72.

28. Fuchs TA, Brill A, Duerschmied D, Schatzberg D, Monestier M, Myers DD, et al. Extracellular DNA traps promote thrombosis. Proc Natl Acad Sci U S A. 2010 Sep 7;107(36):15880–5.

29. Ammollo CT, Semeraro F, Xu J, Esmon NL, Esmon CT. Extracellular histones increase plasma thrombin generation by impairing thrombomodulin-dependent protein C activation. J Thromb Haemost JTH. 2011 Sep;9(9):1795–803.

30. Grover SP, Mackman N. Neutrophils, NETs, and immunothrombosis. Blood. 2018 Sep 27;132(13):1360–1.

31. Deitch EA. Animal models of sepsis and shock: a review and lessons learned. Shock Augusta Ga. 1998 Jan;9(1):1–11.

32. Maloney SK, Fuller A, Mitchell D, Gordon C, Overton JM. Translating animal model research: does it matter that our rodents are cold? Physiol Bethesda Md. 2014 Nov;29(6):413–20.

33. Fu Y, Glaros T, Zhu M, Wang P, Wu Z, Tyson JJ, et al. Network topologies and dynamics leading to endotoxin tolerance and priming in innate immune cells. PLoS Comput Biol. 2012;8(5):e1002526.

